# Cocaine taking and craving produce distinct transcriptional profiles in dopamine neurons

**DOI:** 10.1101/2024.10.11.617923

**Authors:** Tate A. Pollock, Alexander V. Margetts, Samara J. Vilca, Luis M. Tuesta

## Abstract

Dopamine (DA) signaling plays an essential role in reward valence attribution and in encoding the reinforcing properties of natural and artificial rewards. The adaptive responses from midbrain dopamine neurons to artificial rewards such as drugs of abuse are therefore important for understanding the development of substance use disorders. Drug-induced changes in gene expression are one such adaptation that can determine the activity of dopamine signaling in projection regions of the brain reward system. One of the major challenges to obtaining this understanding involves the complex cellular makeup of the brain, where each neuron population can be defined by a distinct transcriptional profile. To bridge this gap, we have adapted a virus-based method for labeling and capture of dopamine nuclei, coupled with nuclear RNA-sequencing, to study the transcriptional adaptations, specifically, of dopamine neurons in the ventral tegmental area (VTA) during cocaine taking and cocaine craving, using a mouse model of cocaine intravenous self-administration (IVSA). Our results show significant changes in gene expression across non-drug operant training, cocaine taking, and cocaine craving, highlighted by an enrichment of repressive epigenetic modifying enzyme gene expression during cocaine craving. Immunohistochemical validation further revealed an increase of H3K9me3 deposition in DA neurons during cocaine craving. These results demonstrate that cocaine-induced transcriptional adaptations in dopamine neurons vary by phase of self-administration and underscore the utility of this approach for identifying relevant phase-specific molecular targets to study the behavioral course of substance use disorders.

## INTRODUCTION

Cocaine use and cocaine-related overdose events have more than tripled over the past two decades [1, 2]. Despite this, there is a shortage of FDA-approved therapeutics for treating cocaine use disorder (CUD) [3-5]. CUD is marked by dysregulated cocaine use and a recurrent cycle of intake, abstinence, and relapse [6, 7]. Therefore, understanding the underlying mechanism of CUD is essential for identifying novel targets to reduce the craving associated with cocaine abstinence, and ultimately lowering relapse rates and overdose events [8-11]. As opposed to natural rewards which engage dopamine (DA) reward circuitry with the greater goal of promoting evolutionary fitness, artificial rewards such as cocaine engage DA circuitry more potently and more persistently than natural rewards, resulting in maladaptive changes in DA signaling that can shift recreational use of a drug toward dependent, and compulsive use [12-15]. Indeed, DA neurons, though constituting less than 1% of central nervous system neurons, play a pivotal role in driving essential neurocognitive processes such as reward-based motivational learning and control of fine motor function [14-16].

While DA neurons from the ventral tegmental are (VTA) represent a primary cellular substrate for drug-induced dopamine release, there are numerous challenges to studying the molecular profile of this population in a cell type-specific manner [6, 13-17]. The cellular heterogeneity of the VTA combined with its small population of DA neurons are two primary obstacles in untangling cell-specific molecular profiles. As such, performing the commonly used technique of bulk RNA-sequencing on VTA tissue may impair the association of drug or behavior-induced gene expression changes specifically to DA neurons [18-20]. Additionally, while novel genomics techniques such as single-cell RNA sequencing have provided unprecedented cellular resolution by generating parallel gene expression profiles of individuals cells, application of these techniques in drug addiction models may be somewhat limited due to the depth of coverage necessary to detect treatment-induced transcriptional changes among lowly expressed genes such as transcription factors and epigenetic enzymes, which are known to drive short- and long-term transcriptional adaptations [21-24].

To overcome these hurdles, we have optimized a method to capture, and sequence VTA DA nuclei extracted from dopamine transporter (DAT)-Cre mice before, during, and after cocaine intravenous self-administration (IVSA) using a Cre-inducible nuclear tag to selectively label dopaminergic nuclei [25-28]. IVSA is a clinically relevant behavioral addiction model as it most closely resembles the drug-taking patterns in humans, in which animals titrate their intake to obtain the maximally rewarding effects of the drug, but before encountering its aversive effects [29-32]. IVSA can be separated into phases that are characterized by drug-taking and drug-craving, consistent with the human drug-taking phase that is followed by a period of abstinence, craving, and in many cases, relapse. The IVSA model mirrors drug-taking in the form of daily sessions where the animal performs and operant task (lever pressing) to earn drug infusions, followed by a period of forced abstinence (craving) where the animal has no contact with the drug-taking environment (operant chamber), and drug-seeking following reintroduction of the drug-experienced animal to the operant chamber [29-31, 33]. Among these phases, understanding the molecular mechanisms underlying drug craving is vital as many individuals suffering with substance use disorders (SUDs) that achieve some level of drug abstinence will eventually relapse, in part due to pervasive incubation of craving that can drive drug-seeking behaviors [8-11].

In this study, we performed cell specific labeling of dopaminergic nuclei in the VTA, followed by cocaine IVSA, nuclear extraction, purification, RNA-sequencing, and computational analyses. We show that cocaine IVSA can drive gene expression profiles in VTA DA neurons that are specific to each phase of the protocol. Indeed, we find enrichment of transcriptionally repressive epigenetic modifying enzymes (G9a, Atf7ip, and Setdb1) during cocaine craving that may shed light into the transcriptional adaptations occurring in DA neurons, as it pertains to relapse risk. To our knowledge, this is the first study combining an *in vivo* nuclear labeling and capture technique with a complex behavioral paradigm to characterize the transcriptome of VTA DA neurons before, during, and after cocaine administration. Importantly, the method detailed herein can be applied to study transcriptional adaptations of genetically defined neurons to other drugs of abuse as well as various other preclinical indications necessitating cell-specific transcriptional profiling of high coverage.

## MATERIALS AND METHODS

### Animals

Male heterozygous B6.SJL-Slc6a3^tm1.1(cre)Bkmm^/J (DAT-Cre; 8-12 weeks old, ∼25-30 g; Jackson Laboratories, Bar Harbor, ME; SN: 006660) and C57BL/6J mice (8-12 weeks old, ∼25-30 g; Jackson Laboratories, Bar Harbor, ME; SN: 000664) were housed in the animal facilities at the University of Miami Miller School of Medicine. Mice were maintained on a 12:12 h light/dark cycle (0600 hours lights on; 1800 hours lights off) and housed three to five per cage. Food and water were provided *ad libitum*. Mice representing each experimental group were evenly distributed among testing sessions. All animals were kept in accordance with National Institutes of Health (NIH) guidelines in accredited facilities of the Association for Assessment and Accreditation of Laboratory Animal Care (AAALAC). All experimental protocols were approved by the Institutional Animal Care and Use Committee (IACUC) at the University of Miami Miller School of Medicine. Whenever possible, the experimenter was blind to the experimental and/or treatment groups.

### Drugs

For self-administration experiments in mice, cocaine hydrochloride (NIDA Drug Supply Program, 96 Research Triangle Park, NC, USA) was dissolved in 0.9% sterile saline.

### Stereotaxic surgery

Mice were anesthetized with an isoflurane (1-3%)/ oxygen vapor mixture and were mounted at a “flat skull” position in a stereotaxic frame (Kopf Instruments, Tujunga, CA). Using aseptic technique, Bregma was exposed by making a 5 mm longitudinal incision on the skin overlying the skull. Two small circular trepanations were drilled in the skull to expose the dura superior to the VTA. Bilateral injections (0.375 µL each at 0.2 µL/min) were made using the following coordinates: VTA, anterior-posterior (AP): -2.95 mm from Bregma; medial-lateral (ML): +/- 0.5 mm from midline; dorsal-ventral (DV): -4.2 mm from dura. The 30-gauge needle was left in place for 5 minutes before retracting to ensure proper AAV5-DIO-KASH-HA viral dispersion.

### Jugular catheterization

Jugular catheterization was performed as previously described [34]. Briefly, mice were anesthetized with an isoflurane (1–3%)/oxygen vapor mixture and prepared with indwelling jugular catheters. Briefly, the catheters consisted of a 6.5-cm length of Silastic tubing fitted to guide cannulas (PlasticsOne, Protech International Inc., Boerne, TX, USA) bent at a curved right angle and encased in dental acrylic and silicone. Catheter tubing was subcutaneously passed from the animal’s back toward the right jugular vein. 1 cm of the catheter tip was inserted into the vein and secured with surgical silk sutures. Mice were administered Meloxicam (5 mg/kg) subcutaneously before surgery and 24 hours post-surgery. Catheters were flushed daily with sterile saline solution (0.9% w/v) containing heparin (10–60 USP units/mL) beginning 48 hours after surgery. Animals were allowed 3-5 days to recover before commencing intravenous cocaine self-administration. Catheter integrity was tested with the ultra-short-acting barbiturate anesthetic Brevital (methohexital sodium, Eli Lilly, Indianapolis, IN, USA).

### Operant food training and cocaine intravenous self-administration

The intravenous self-administration (IVSA) procedure measures the reinforcing properties of a drug. Mice self-administered intravenous cocaine infusions in daily 1-hour sessions, using a reinforcement schedule of FR5TO20, where meeting a fixed ratio (FR) of 5 consecutive active lever presses resulted in delivery of an IV infusion and presentation of a 20s cue light, which coincided with a 20s time-out (TO) period during which active lever presses did not count toward delivery of a reward. Prior to IVSA, mice underwent 7 consecutive days of food training during which the FR schedule increased from FR1 to FR5, provided the mouse self-administered >30 food pellets per session at a given FR. Following food training, mice underwent jugular catheter implantation and resumed food training. After confirming maintained lever pressing behaviors post-surgery, mice were divided into “Food Trained” or “Cocaine IVSA” groups based on food rewards earned to avoid baseline differences in operant performance from biasing the behavioral and transcriptional results (**Supplemental Figure 1**). The brains of the Food Trained mice were then collected for molecular analyses.

IVSA mice proceeded to complete 5 consecutive days of cocaine acquisition using a dose of 0.3 mg/kg/inf (0.032 mL infusion) at FR5TO20. This was followed by a maintenance phase where mice self-administered cocaine (1.0 mg/kg/inf) at FR5TO20 for 10 consecutive days, totaling 15 consecutive days of cocaine IVSA. Catheters were flushed daily with a heparinized saline solution (10 U/mL, 0.05 mL) prior to, and immediately following each IVSA session. Mice that failed to show stable cocaine responding (>25% variation in intake across 3 consecutive days), that failed to meet threshold for intake (< 6 infusions per 1 hour session), or displayed compromised catheter patency, were excluded from analysis.

### Forced home-cage abstinence and cocaine-seeking

Following maintenance, “Cocaine IVSA” mice were divided into “Cocaine-Taking”, or “Cocaine-Craving” groups based on the number of drug rewards earned, thus ensuring no significant differences in cocaine taking behavior between groups. The brains of the “Cocaine-Taking” mice were then collected for molecular analyses, and the remaining “Cocaine-Craving” mice underwent forced home-cage abstinence for 21 days, where they were placed in their home cages without access to cocaine and thus without access to environmental cues associated with the drug-taking environment [10, 11]. We chose the 21-day timepoint for craving, as reintroducing the animals to the IVSA chamber after this time resulted in robust active lever pressing (cocaine-seeking), despite the absence of any drug reward—akin to a cocaine reinstatement session (**Fig. 1F**). It should be noted that the brains of the “Cocaine-Craving” mice were collected after 21 days of home-cage abstinence and no cocaine-seeking session was conducted on these animals. During brain collections at the end of each phase, all mice were euthanized by standard methods (isoflurane followed by transcardial perfusion for histological analyses, and decapitation for molecular analyses).

**Figure 1.**
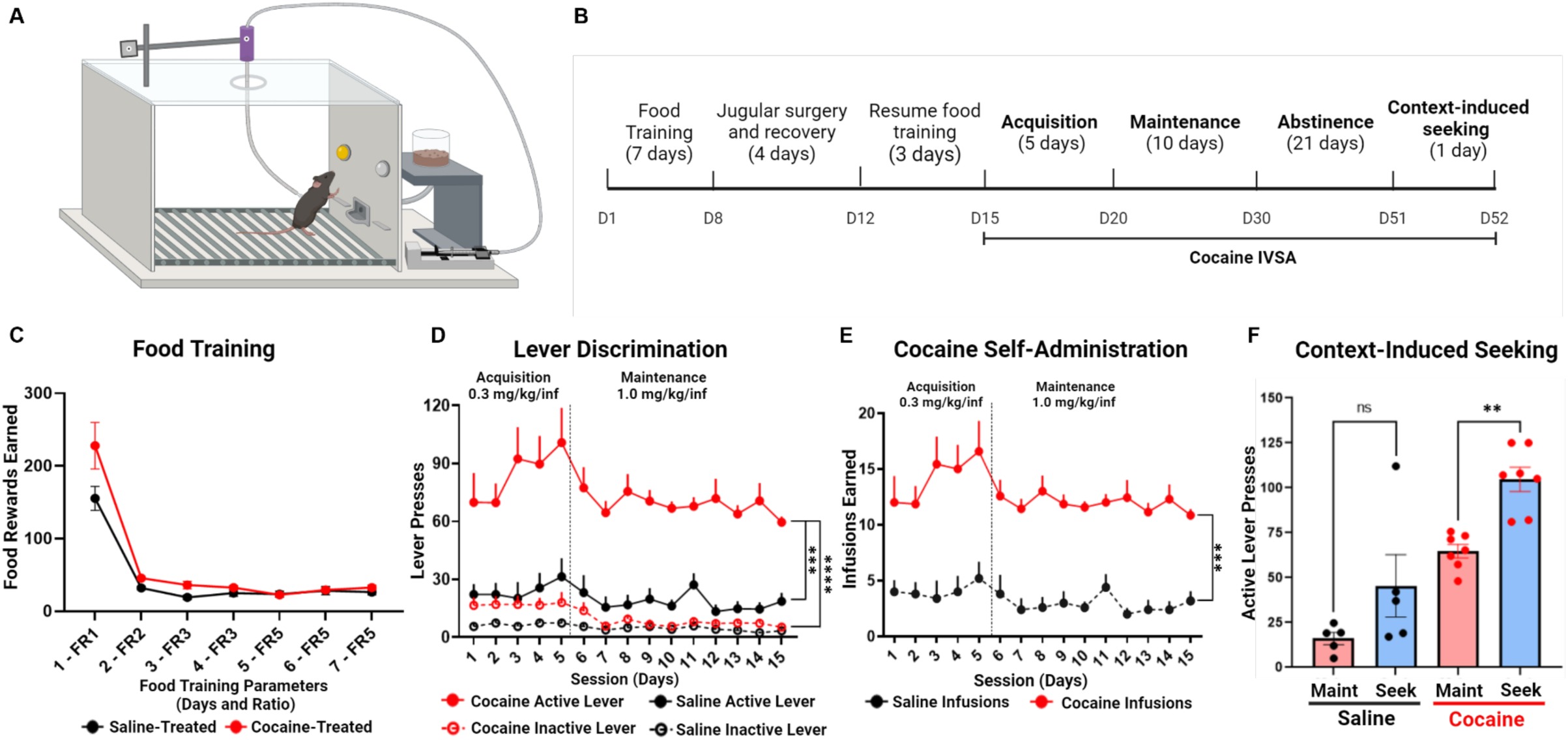
Mice acquire and maintain stable cocaine intravenous self-administration. **A)** The operant chamber apparatus for cocaine nous self-administration (IVSA). **B)** Experimental timeline. **C)** Average food rewards earned daily during 7 days of food training n treatment groups. Two-way RM ANOVA (n=5-7, Cocaine vs Saline Food Reward, ns). FR = fixed ratio. **D)** Daily active and e lever presses between saline- and cocaine-treated mice throughout 5 days of acquisition and 10 days of maintenance. Two-way OVA with Bonferroni’s post-hoc test (n=5-7, Cocaine Active vs Inactive Lever, ****p < 0.0001; Cocaine vs Saline Active Lever, 0.0003). **E)** Daily infusions earned between saline- and cocaine-treated mice. Two-way RM ANOVA with Bonferroni’s post-hoc test , Saline Infusions vs Cocaine Infusions, ***p = 0.0001). **F)** Merged graph of final 3-day average of active lever presses during nance (**Maint**) compared to context-induced seeking (**Seek**) in both saline- and cocaine-treated mice. Two-tailed paired t test (n=5, Maint vs Seek, ns; n=7, Cocaine Maint vs Seek, **p = 0.0013). Data are represented as mean ± SEM. Created with *BioRender*.

### Isolation of dopamine nuclei

Mice were anesthetized with isoflurane, decapitated, and the brains were removed. Using a brain block, the brains were sliced coronally to reveal the ventral tegmental area (VTA). Bilateral tissue punches from the VTA of 2 mice were combined for each sample and transported in Hibernate A Medium (Gibco, A1247501). Nuclei were mechanically and enzymatically isolated using the Nuclei Extraction Buffer (Miltenyi Biotec, 130-128-024) following manufacturer’s instructions. Suspensions were filtered through 100 µM and 30 µM filters after repeated centrifugations at 300 x g for 5 minutes at 4^°^C. The resulting nuclei were resuspended in Wash Buffer (1 mL 1M HEPES pH 7.5, 1.5 mL 5M NaCl, 12.5 µL 2M spermidine, 1 Roche c0mplete Protease Inhibitor EDTA-free tablet with ddH2O to 50 mL) [35] with 10% DMSO and slow frozen in a Mr. Frosty (ThermoFisher, 5100-0001) at -80° C and stored until processed for fluorescently activated nuclear sorting (FANS).

### Nuclear immunostaining and fluorescent activated nuclear sorting (FANS)

Frozen nuclear suspensions were submerged in a 20mL beaker containing ddH20 at RT and allowed to thaw. Nuclei were centrifuged at 1000 x g for 5 minutes at 4^°^C and incubated in 1 mL blocking solution (2.5 mM MgCl^2^, 1% BSA, 0.2 U/µL RNAsin in PBS) for 30 minutes at 4^°^C. After blocking, nuclei were incubated with Alexa Fluor 647 Conjugated HA-Tag (6E2) Mouse mAb (1:50, Cell Signaling Technology, #3444) in blocking solution for 1 hour at 4°C on a tube rotator. After antibody incubation, 64 µL NucBlue Fixed Cell ReadyProbes Reagent (DAPI) (Invitrogen, #37606) in 500 µL blocking solution was added to nuclei for 20 min at 4°C. The immunostained nuclei were transferred into filtered FACS tubes and sorted with a CytoFLEX SRT (Beckman Coulter) equipped with Violet (405 nm), Blue (488 nm), Yellow/Green (561 nm), and Red (640 nm) lasers at the Flow Cytometry Shared Resource (FCSR) of the Sylvester Comprehensive Cancer Center at the University of Miami. All samples were sorted based on the nuclear size, complexity and a positive AF-647 and DAPI signal, and the resulting isolated populations were either AF-647^+^(HA^+^)/DAPI^+^ (DA nuclei) or AF-647^-^(HA^-^)/DAPI^+^ (nDA).

### Brain perfusion and fixation

Mice were anesthetized with isoflurane and perfused through the ascending aorta with PBS pH 7.4 (Gibco, 10010023) plus heparin (7,500 USP units), followed by fixation with 4% paraformaldehyde in PBS. Brains were collected, postfixed overnight in 4% paraformaldehyde, and transferred to 30% sucrose with 0.05% sodium azide (S2002, Sigma-Aldrich, St. Louis, MO, USA) in PBS for 72 hours. All brains were cut into 25-30 µm coronal free-floating sections on a Leica CM1900 cryostat and placed into 12-well plates containing PBS with 0.02% sodium azide at 4°C until processing for immunohistochemistry.

### Immunohistochemistry

Floating sections were processed for fluorescent immunostaining of dopamine neurons. Sections were rinsed in PBS, then blocked for 1 hour in Blocking Buffer (10% normal donkey serum (017-000-121, Jackson ImmunoResearch), 0.2% Triton X-100 (T8787, Sigma), and PBS). Sections were then incubated with primary antibodies diluted in blocking buffer overnight at 4°C. The primary antibodies used were: mouse anti-Th (1:500, SC-25269, Santa Crus Biotechnology), rabbit anti-G9a (1:500, GTX129153, GeneTex), rabbit anti-SETDB1 (1:250, 11231-1-AP, Proteintech Group), and rabbit anti-H3K9me3 (1:500, ab8898, Abcam). On day 2, sections were washed in PBS three times for 5 minutes each, then incubated with the following secondary antibodies: Alexa 488 Donkey anti-Mouse (1:500, A21202, Invitrogen) and Alexa 568 Donkey anti-rabbit (1:500, A10042, Invitrogen). Sections were incubated with secondary antibodies in PBS with 2% normal donkey serum for 2 hours at room temperature in the dark. Sections were then rinsed in PBS three times for 5 minutes each, mounted on slides with ProLong Diamond Antifade Mountant with DAPI (Invitrogen, P36962) and cover-slipped. Fluorescent images were acquired on an ECHO Revolve microscope using 4X, 10X, and 20X objectives and saved as both grayscale and pseudo-colored .tiff images. All antibodies used have been previously validated for the intended applications, as per manufacturer. In 12 animals, the immunolabeling experiment was successfully repeated for all representative images of qualitative data.

### Corrected total cell fluorescence (CTCF) immunohistochemistry quantification

To identify Th+ neurons and assess histone modification levels, brain sections were prepared and stained for tyrosine hydroxylase (Th) and histone H3 lysine 9 trimethylation (H3K9me3). Images were acquired at 20X magnification using a fluorescence microscope. Each of the three treatment groups (Food Trained, Cocaine-Taking, & Cocaine-Craving) was composed of four mice. Four brain sections were imaged per mouse, 25 Th+ cells were identified and analyzed per section. Using ImageJ software, regions of interest (ROIs) were manually drawn around the nuclei of each of the 25 Th+ cells in the section. For each ROI, the integrated density (the product of area and mean fluorescence intensity) was measured for H3K9me3 staining. Background fluorescence was measured by selecting 7 areas devoid of specific staining in each section. The mean fluorescence intensity in these areas was calculated and averaged to determine the background level for each section. This average background fluorescence was subtracted from the fluorescence intensity of each cell to obtain the corrected total cell fluorescence (CTCF). The CTCF for each cell was averaged to obtain the mean CTCF per cell per section. Additionally, the total CTCF for all 25 cells was calculated in each section to determine the total section fluorescence. These calculations allowed for the comparison of H3K9me3 levels across all groups. Statistical analyses were performed to compare the mean and total CTCF across the groups. One-way ANOVA with multiple comparisons was used to determine significant differences between groups and a p-value < 0.05 was considered statistically significant.

### RNA-sequencing

Total RNA was directly isolated from sorted nuclei using the Qiagen AllPrep DNA/RNA Mini Kit (Qiagen, 80204). Briefly, nuclei were sorted into ∼700 µL RLT plus buffer (Qiagen) for the extraction and purification of RNA. DNA was eliminated via column exclusion and RNA was purified following manufacturer’s instructions. RNA-sequencing libraries were prepared using the NEBNext Single Cell/Low Input RNA Library Prep Kit for Illumina (New England BioLabs, E6420S) after normalizing RNA input. Paired-end 150 bp sequencing was performed on a NovaSeq6000 sequencer (Illumina) by the University of Miami Center John P. Hussman Institute for Human Genomics sequencing core facility, targeting 30 million reads per sample. Raw RNA-seq datasets were first trimmed using Trimgalore (v.0.6.7) and cutadapt (v.1.18). Illumina adaptor sequences were removed, and the leading and tailing low-quality base-pairs were trimmed following default parameters. Next, the paired-end reads were mapped to the mm10 mouse genome using STAR (v.2.7.10a) with the following parameters: --outSAMtype BAM SortedByCoordinate –outSAMunmapped Within –outFilterType BySJout –outSAMattributes NH HI AS NM MD XS –outFilterMultimapNmax 20 –outFilterMismatchNoverLmax 0.3 –quantMode TranscriptomeSAM GeneCounts. The resulting bam files were then passed through StringTie (v.2.1.5) to assemble sequence alignments into an estimated transcript and gene count abundance given the NCBI RefSeq GRCm38 (mm10) transcriptome assembly.

### Differential Gene Expression Analysis

The R/Bioconductor DESeq2 package (v.1.38.3) [36] was used to detect the differentially expressed genes between VTA DA nuclei and all other VTA nuclei, as well as VTA DA nuclei throughout different phases of cocaine IVSA. Following filtering for low count genes, as determined by DESeq2, only genes with a False Discovery Rate (FDR) adjusted p-value (padj) < 0.05 were considered significantly differentially expressed. In the case where biological replicates showed large variability indicating outliers a supervised removal of such replicates from each group was conducted. Heatmaps were generated using the R/Bioconductor package pheatmap (v.1.0.12) of log transformed normalized counts from merged lists of significantly differentially expressed genes using the following filters: padj < 0.05, log2foldchange (L2FC) ≥ 1.5 in a pairwise comparison, and a log foldchange standard error (lfcSE) ≤ 1. Heatmaps were clustered based on correlation using the “ward.D” method. Venn diagrams were generated by filtering significant results lists of genes (padj < 0.05) for up and down-regulated genes based on log2foldchange values greater than 0 and less than 0 for each pairwise comparison. All the other plots were generated using ggplot2 package (v.3.4.2).

### Functional Enrichment Analysis

The enrichGO function from the R/Bioconductor clusterProfiler package (v.4.6.2) was used to perform gene ontology (GO) enrichment analysis. Only significantly differentially expressed genes with an adjusted p value ≤ 0.05 and a log-fold change standard error ≤ 1.5 were included, while also removing genes that did not map to Entrez identifiers. Resulting GO terms and pathways with an FDR corrected q-value < 0.10 and padj < 0.05 were considered after using a custom background from all genes that were expressed after DESeq2 adjustment. The associated GO and pathway enrichment plots were generated using the ggplot2 package.

### Statistical analyses

Previous results from our lab [34] and post-hoc power analyses of preliminary data were used to estimate reasonable sample size. The data distribution was assumed to be normal. For self-administration experiments, animals that did not achieve stable levels of cocaine intake (>25% variation in intake across 3 consecutive days) or that earned fewer than 6 cocaine infusions on average across sessions were excluded from data analysis. All data were analyzed by one-way or two-way ANOVAs with multiple comparisons or Bonferroni’s post-hoc test, or paired t-tests using GraphPad Prism software (La Jolla, CA). Significant main or interaction effects were followed by multiple comparison tests. The criterion for significance was set at *p* ≤ 0.05. Results are shown as the mean ± SEM. All statistical analyses used in differential gene expression analyses are generated from the established DESeq2 package [36] with minor modifications to obtain more stringent results.

## RESULTS

### Mice acquire and maintain stable cocaine intravenous self-administration

To establish the mouse cocaine IVSA model, C57BL/6 mice underwent cocaine or saline intravenous self-administration (IVSA) in an operant chamber (Med Associates Inc, Fairfax, VT, USA) (**Fig. 1A**) following the experimental timeline in **Figure 1B**. Mice were assigned to treatment groups (Saline-treated or Cocaine-treated) based on similar average food rewards earned during food training (**Fig. 1C**) (Two-way RM ANOVA; n = 5-7, Cocaine vs Saline Food Rewards, F(1, 10)=4.918, p=0.0509, ns). Cocaine-treated mice demonstrated significant lever discrimination in favor of the active lever over the inactive lever, while saline-treated animals failed to demonstrate a lever preference in the operant task (**Fig. 1D**) (Two-way RM ANOVA; n = 5-7, Cocaine Active vs Inactive Lever Presses, F(1, 12)=66.25, p<0.0001). Additionally, active lever responding was significantly higher in cocaine-treated mice than in saline-treated mice (**Fig. 1D**) (Two-way RM ANOVA; n = 5-7, Cocaine vs Saline Active Lever Presses, F(1,10)=30.04, p=0.0003), as well as number of IV infusions earned per session (**Fig. 1E**) (Two-way RM ANOVA; n = 5-7, Cocaine vs Saline Infusions, F(1,10)=38.49, p=0.0001), highlighting the reinforcing properties of cocaine using this behavioral paradigm. Following the conclusion of cocaine or saline maintenance, mice were subjected to 21 consecutive days of forced home-cage abstinence. Following abstinence, mice completed a 1-hour cocaine- or saline-seeking session (as described in the methods) in which cocaine-treated mice responded on the active lever at a significantly higher rate than their respective response rates during maintenance (**Fig. 1F**) (Two-tailed paired t test; n = 7, Cocaine Maint vs Seek, p=0.0013). No such significant effect was observed in saline-treated mice (**Fig. 1F**) (Two-tailed paired t test; n = 5, Saline Maint vs Seek, ns), suggesting that incubation of cocaine craving over 21 consecutive days of forced home-cage abstinence can result in relapse-like operant responding when the animal is reintroduced to the drug-taking environment.

### Cell-specific nuclear labeling reveals enrichment of dopaminergic transcriptome

To obtain a pure population of VTA DA neurons, we employed a cell type-specific nuclear labeling and capture technique previously established by Tuesta et al. [25]. Briefly, this method (**Fig. 2A**) involves the stereotaxic introduction of a Cre-inducible adeno-associated virus (AAV) vector encoding the nuclear envelope protein KASH with a hemagglutinin (HA) tag (KASH-HA) (**Fig. 2B**) into the VTA of dopamine transporter (DAT)-Cre^+/-^ mice, thereby selectively labeling midbrain DA nuclei from which intact chromatin and nascent RNA can be isolated [25]. In its present application, RNA-Sequencing was conducted on RNA obtained from isolated and sorted HA^+^ (DA) and HA^-^ (non-DA) nuclear populations.

**Figure 2.**
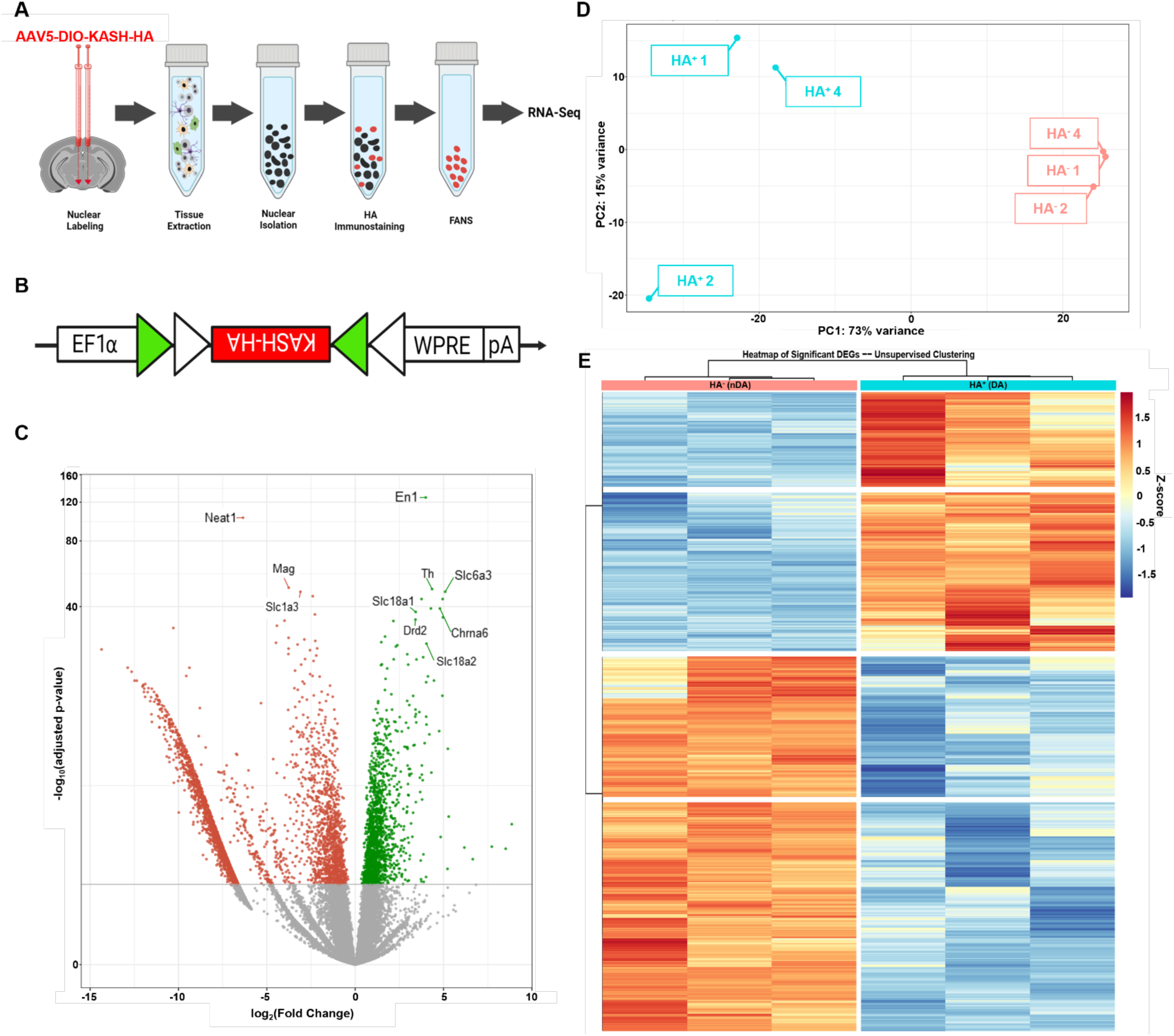
Dopamine nuclei can be labeled and captured with viral injections and RNA-sequenced. **A)** An overall outline for nuclear ling and capture from viral injection to RNA-Seq. **B)** The Cre-inducible adeno-associated viral vector encoding KASH with an HA tag to label DA nuclei. **C)** Volcano plot of HA+ vs HA-differentially expressed genes (DEGs) (Green: significantly upregulated in HA+ ei, Red: significantly downregulated in HA-nuclei, Grey: not significantly up or down-regulated; padj < 0.05, lfc > 0 or lfc < 0; Genes portance marked with lines and labels). **D)** Principal component analysis of variance between HA+ nuclei (cyan) and HA-nuclei mon). **E)** Unsupervised clustering heatmap of significant DEGs (padj <0.05, L2FC ≥ 1.5 and L2FC ≤ 1.5, lfcSE ≤ 1) between HA+ ei (cyan) and HA-nuclei (salmon). n = 3. Panels A and B created with *Biorender*.

Differential gene expression analyses reveal that out of the 26,336 genes expressed (a nonzero read count) in HA^+^ and HA^-^ nuclei, 113 genes exhibited at least a 2.5-log2fold enrichment in HA^+^ cells, including DA identity genes such as Th, Drd2, Slc6a3 (Dat), and Slc18a2 (Vmat2) (**Fig. 2C, Supplemental Table 1**). Conversely, 1,742 genes were significantly depleted (padj < 0.05, log2foldchange < -2.5) in HA^+^ compared to HA^-^ nuclei, including oligodendrocyte and astrocyte markers such as Mag [37] and Slc1a3 [38], respectively (**Fig. 2C, Supplemental Table 1**). Both principal component analysis (PCA) (**Fig. 2D**) and unsupervised clustering analyses (**Fig. 2E**) highlight the contrast between the overall transcriptional profiles of HA^+^ and HA^-^ populations.

### VTA DA neurons exhibit distinct transcriptional profiles during Cocaine-Taking and Cocaine-Craving

We next asked whether cocaine IVSA induced gene expression changes in VTA DA neurons, and if these changes varied by phase of self-administration (**Fig. 3A**). To this end, we profiled the gene expression profiles of DAT-Cre^+/-^ mice undergoing the cocaine IVSA paradigm, as described in **Fig. 3A**. Indeed, PCA revealed phase-specific clustering of gene expression from samples during non-drug operant training (Food Trained, n = 3), cocaine maintenance (Cocaine-Taking, n = 3), and home-cage abstinence (Cocaine-Craving, n = 4), wherein the Cocaine-Taking cohort showed the most variance from the others, while the Cocaine-Craving cohort clustered around the tight Food Trained cluster (**Fig. 3B**). Further differential gene expression analyses, utilizing DESeq2 [36] to conduct pairwise comparisons, revealed 1,301 differentially expressed genes (DEGs) (padj < 0.05) detected in “Food Trained vs Cocaine-Taking” and 1,050 DEGs (padj < 0.05) detected in “Cocaine-Taking vs Cocaine-Craving,” while there were overall only 247 DEGs (padj < 0.05) found in “Food Trained vs Cocaine-Craving” (**Fig. 3C**). Dividing the total DEGs further into upregulated and downregulated genes revealed that pairwise comparisons of VTA DA nuclei involving Cocaine-Taking are distinctive from one another, while DEGs found in “Food Trained vs Cocaine-Craving” share large overlap with those detected in both Cocaine-Taking comparisons (**Fig. 3D-E**).

**Figure 3.**
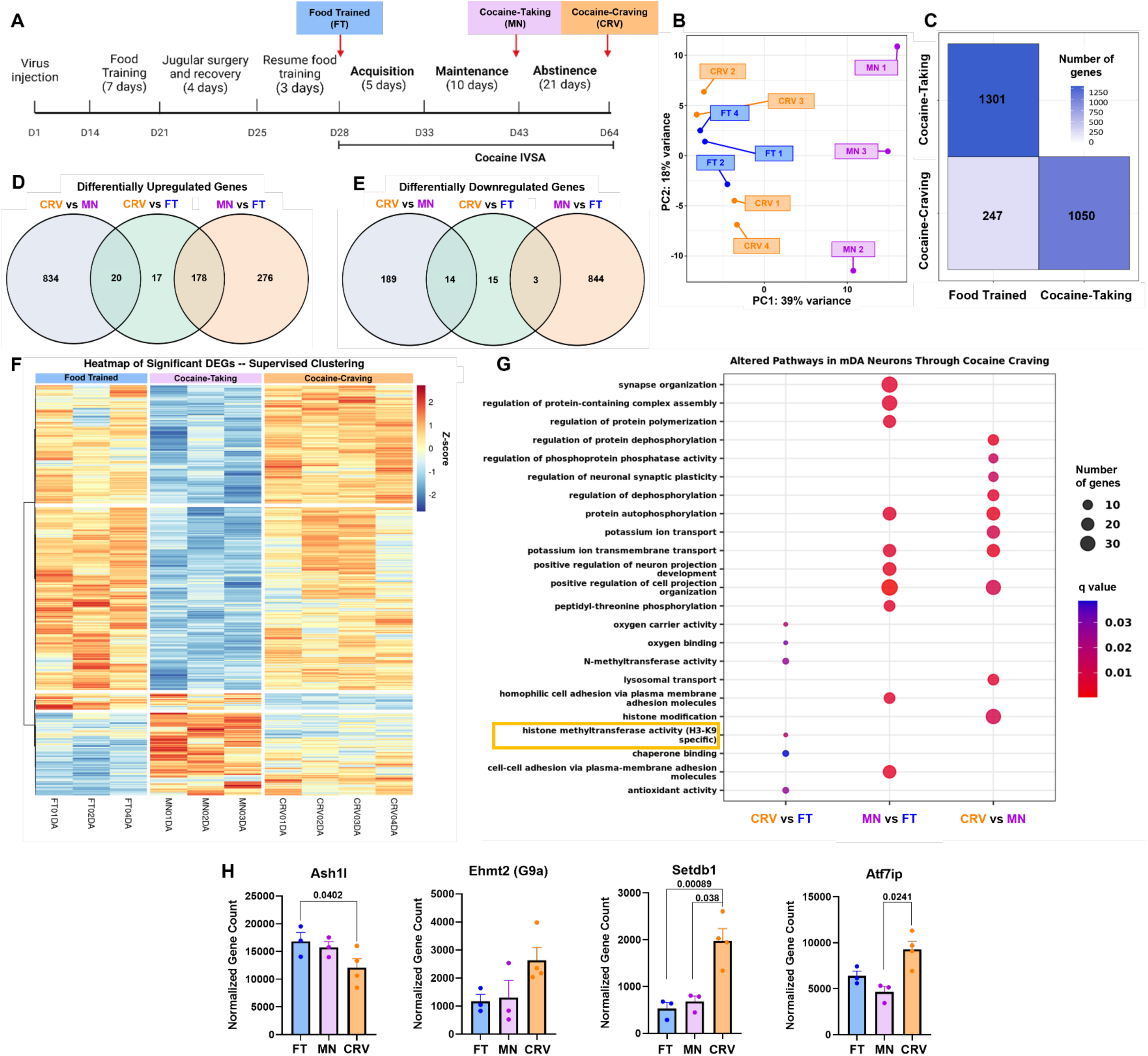
RNA-Sequencing reveals DA-specific differentially expressed genes throughout cocaine intravenous self-administration. **A)** tal timeline for cocaine intravenous self-administration (CIVSA) with KASH-HA injections. Red arrows indicate tissue extraction days t labels. Food Trained, n=3 (FT, blue); Cocaine-Taking, n=3 (MN, purple); and Cocaine-Craving, n=3 (CRV, orange). **B)** Principal t analysis of variance in RNA-seq between CIVSA phase-specific VTA DA nuclei. **C)** Differentially expressed genes (DEGs) in pairwise ns of CIVSA phase-specific VTA DA nuclei (padj < 0.05). **D)** All upregulated and overlapping genes found in pairwise comparisons of ase-specific VTA DA nuclei (padj < 0.05). Upregulated gene counts correspond to the first cohort listed in each pairwise comparison. **E)** gulated and overlapping genes found in pairwise comparisons of CIVSA phase-specific VTA DA nuclei (padj < 0.05). Downregulated ts correspond to the first cohort listed in each pairwise comparison, L2FC < 0. **F)** Heatmap of top 426 significant DEGs in HA^+^ nuclei ood Trained, Cocaine-Taking, and Cocaine-Craving mice during CIVSA (padj < 0.05, L2FC ≥ 1.5 or L2FC ≤ -1.5, lfcSE ≤ 1). **G)** Gene GO) pathway analysis conducted using DEGs (padj < 0.05, lfcSE ≤ 1.5) found between pairwise comparisons of phases of CIVSA (padj lue < 0.10, Selected pathways shown for each comparison). **H)** Normalized counts of genes involved in transcriptional methyltransferase between phases of CIVSA (values represent significant adjusted p-values between pairwise comparisons using DESeq2).

To obtain a visual representation of the directionality of gene expression changes, we next generated a heatmap of top differentially expressed genes by phase of IVSA. Consistent with **Fig. 3C-E**, most gene expression changes occurred during Cocaine-Taking. More specifically, cocaine IVSA induced transcriptional repression in DA neurons that largely returned to baseline following 21 days of abstinence in Cocaine-Craving animals (**Fig. 3F**). Indeed, this finding was buttressed by gene ontology (GO) pathway analysis where the gene expression patterns of the Cocaine-Craving cohort exhibit many compensatory adaptations restoring most genes to Food Trained levels, as numerous significant GO pathways were identified in pairwise comparisons of both “Cocaine-Taking vs Cocaine-Craving” and “Cocaine-Taking vs Food Trained” conditions, while only 6 significant GO pathways were identified between “Food Trained and Cocaine-Craving” conditions (**Fig. 3G**). For example, we identified one significantly enriched GO pathway in “Cocaine-Taking vs Cocaine-Craving” named “histone modifications” (**Fig. 3G**) which, like (**Fig. 3F**), illustrates how the changes made during cocaine maintenance normalized in the absence of the drug. However, a separate GO pathway named “histone methyltransferase activity (H3-K9 specific)” was significantly enriched in “Food Trained vs Cocaine-Craving” suggesting that while many cocaine-induced transcriptional changes normalized, there may still be dysregulation of differentially expressed genes in the absence of the drug (**Fig. 3G**).

This finding led to the identification of individual genes involved in H3K9 methylation. Several such genes involved in transcriptional regulation (Ash1l, Ehmt2, Setdb1, & Atf7ip) via post-translational modifications (PTMs) of H3K9 were found to be most differentially expressed during Cocaine-Craving (**Fig. 3H**), suggesting that cocaine craving may also drive differential deposition of H3K9 methylation. Additionally, while cocaine administration drives much of the DEGs, these data suggest that Cocaine-Craving animals do not fully restore the alterations made to genes in VTA DA neurons during Cocaine-Taking, resulting in enrichment of H3K9 methyltransferases.

### H3K9me3 is enriched in DA neurons of Cocaine-Craving mice

**Fig. 3H** suggests the differential expression of the methyltransferases Ahs1l, Ehmt2, Setdb1, and Atf7ip in the Cocaine-Craving cohort, and it is known that one of their products, H3K9me3, is a transcriptionally repressive histone mark with many downstream effects [39]. Therefore, we used H3K9me3 as a surrogate marker of this methyltransferase activity. To this end, we performed immunostaining of H3K9me3 and the dopaminergic marker tyrosine hydroxylase (Th) on VTA-containing midbrain sections from separate cohorts of C57BL/6 mice that completed the behavioral paradigm described in **Fig. 3A**. A schematic of this brain region is shown, accompanied by a merged 4X bilateral and 10X unilateral representative image (**Fig. 4A**). Quantifications were taken at 20X and representative images from the 3 treatment cohorts are shown (**Fig. 4B**). The average corrected total cell-fluorescence (CTCF) per section was measured and revealed significantly higher average fluorescence intensity of H3K9me3 in Th^+^ neurons from the Cocaine-Craving cohort (**Fig. 4C**) (One-way ANOVA with Tukey’s multiple comparisons; n = 16, Food Trained vs Cocaine-Craving, p<0.0001; Cocaine-Taking vs Cocaine-Craving, p<0.0001; Food Trained vs Cocaine-Taking, p=0.4385, ns). Additionally, all CTCF measurements were averaged per mouse, with Cocaine-Craving mice displaying significantly higher average cell fluorescence intensity of H3K9me3 in Th^+^ neurons, while no significant difference was observed in the Food Trained or Cocaine-Taking mice (**Fig. 4D**) (One-way ANOVA with Tukey’s multiple comparisons; n=4, Food Trained vs Cocaine-Craving, ***p = 0.0007; Cocaine-Taking vs Cocaine-Craving ***p = 0.0003; Food Trained vs Cocaine-Taking, p = 0.7010, ns).

**Figure 4.**
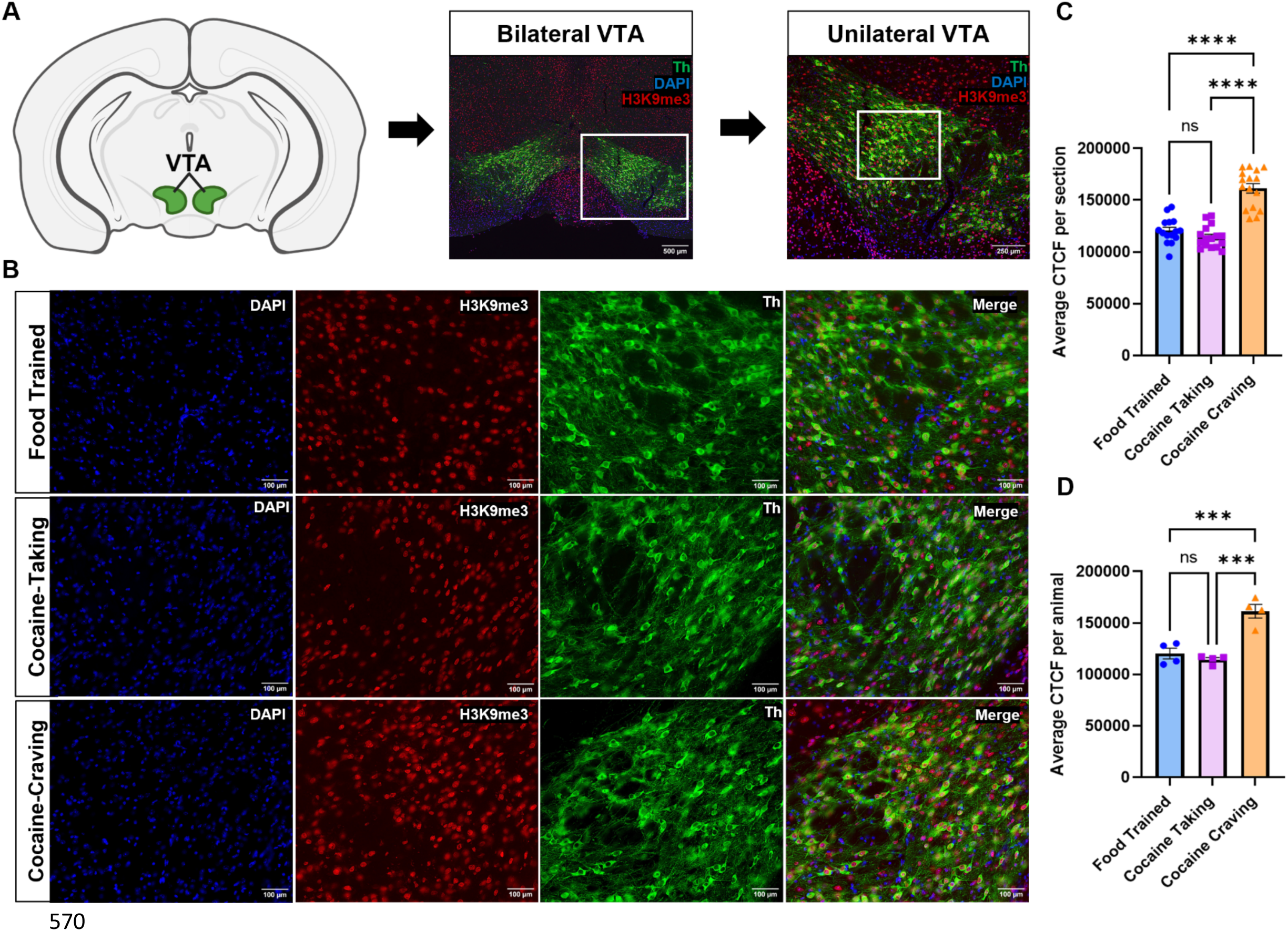
H3K9me3 is enriched in DA neurons of Cocaine-Craving mice. **A)** Graphical representation of the VTA in a coronal mouse brain slice ine hydroxylase (Th, green), histone-3 lysine-9 trimethylation (H3K9me3, red), and DAPI (blue) in a 4x bilateral VTA merged image. 500µm (middle). A 10x unilateral VTA merged image. Scale bar: 250µm (right). **B)** Representative 20x images of the VTA in Food caine-Taking, and Cocaine-Craving mice. Scale bar: 100µm. **C)** Average corrected total cell fluorescence (CTCF) per section between ed, Cocaine-Taking, and Cocaine-Craving groups. n=16; One-way ANOVA with Tukey’s multiple comparisons. Food Trained vs aving, ****p < 0.0001; Cocaine-Taking vs Cocaine-Craving ****p < 0.0001; ns = not significant. **D)** Average CTCF per animal between oups. n=4; One-way ANOVA with Tukey’s multiple comparisons. Food Trained vs Cocaine-Craving, ***p = 0.0007; Cocaine-Taking vs aving ***p = 0.0003; ns = not significant. Data are represented as mean ± SEM.

## DISCUSSION

Dopamine (DA) signaling underpins a variety of neurological and psychiatric disorders, yet our understanding of gene expression patterns within DA neurons in both health and diseased states remain poorly understood [17-19, 40, 41]. Here, we leveraged an established molecular method for cell type-specific labeling and capture of VTA DA neurons to identify gene expression changes associated with cocaine reinforcement and craving. This approach complements previous transcriptional profiling efforts by providing cell type resolution to further contextualize gene expression changes through the prism of dopaminergic signaling.

### VTA DA neurons exhibit transcriptional profiles specific to phase of IVSA

We expected to identify gene expression changes consistent with cocaine use and abstinence in the Cocaine-Taking and Cocaine-Craving cohorts, respectively. Indeed, relative to Food Trained control mice, we found 1301 DEGs in the Cocaine-Taking cohort and 247 DEGs in the cohort of mice undergoing cocaine craving (**Fig. 3 C-E**). Similar to the transcriptional consistency of VTA DA neurons shown with unsupervised DEG clustering (**Fig. 2E**), the unsupervised DEG clustering of these neurons in mice undergoing either Food Training, Cocaine-Taking, or Cocaine-Craving resulted in clustering by phase of self-administration (**Sup Fig. 3**). Unsupervised clustering of biological replicates by treatment suggests that each phase of cocaine intravenous self-administration features a unique dopaminergic transcriptional profile.

Dopamine neurons exhibit adaptive transcriptional changes secondary to cocaine self-administration that largely, but not completely, normalize following a 21-day period of abstinence. When framed in the context of hedonic allostasis [7, 42, 43], the “allostatic” transcriptional profile of Cocaine-Craving mice contains a number of genes that are either persistently dysregulated from the Cocaine-Taking phase or newly dysregulated following cessation of the drug. These maladaptive transcriptional changes may lend insight into the neurobiological mechanisms underlying cocaine craving and relapse [11].

### Alterations to H3K9-specific methyltransferase expression in VTA DA neurons

Epigenetic regulation of transcription plays a central role in the magnitude, duration, and latency of gene expression. GO analysis revealed enrichment of DEGs associated with histone modification, but more specifically, we detected enrichment of DEGs associated with H3K9 methyltransferase activity (**Fig. 3G**). Methylation of H3K9 is known to be transcriptionally repressive. To this end, we identified four epigenetic regulatory genes: absent, small, or homeotic (ASH) 1-like histone lysine methyltransferase (Ash1l), euchromatic histone lysine methyltransferase 2 (Ehmt2; commonly known as G9a), SET domain bifurcated histone lysine methyltransferase 1 (Setdb1), and activating transcription factor 7 interacting protein (Atf7ip). Evidence suggests that these genes and their protein products play various roles throughout the brain reward system, discussed below.

Ash1l is a histone methyltransferase (HMT) with numerous interaction domains that mediates methylation throughout the genome from the di-methylation of H3K36 [44], to working in a DNA repair complex that tri-methylates H3K4 [45]. The crystal structure of Ash1l reveals multiple catalytic domains, with one such domain being the mono-methylation of H3K9 (H3K9me) [46]. H3K9me is considered a prerequisite for the further deposition of H3K9 methylation as performed by G9a and Setdb1 [39]. Our data show that Ash1l is significantly downregulated during Cocaine-Craving, which suggests that further methylation of H3K9 (H3K9me2, H3K9me3) may involve the involvement of additional methyltransferases such as G9a and Setdb1.

Ehmt2 (G9a) is a methyltransferase that primarily mono- and di-methylates histone 3, lysine 9 (H3K9me/ H3K9me2) [47-49]. While G9a has become a gene of interest in the brain reward system, its involvement in dopaminergic signaling is less understood. We found downregulation of G9a in both Food Trained and Cocaine-Taking groups, followed by a trending enrichment of G9a during Cocaine-Craving (**Fig. 3H**). Interestingly, these results mirror trends in the NAc, where G9a acts bidirectionally. In the NAc, cocaine reduces G9a levels during use [50-52], while after weeks of forced home-cage abstinence, a knockdown of G9a mitigates cocaine-primed reinstatement [53]. This suggests that the increased G9a expression we find during abstinence could contribute to drug-seeking behaviors [52, 54]. Moreover, overexpression of G9a in the NAc increases sensitivity to cocaine and reinstates cocaine-seeking in rats [53], underscoring the importance of G9a and its repressive H3K9me2 in reward systems during cocaine administration.

Similar to G9a, we found downregulation of Setdb1 in Food Trained and Cocaine-Taking groups, followed by a significant upregulation during Cocaine-Craving (**Fig. 3H**). Setdb1 is a repressive methyltransferase, depositing H3K9me2 and H3K9me3 in both histone and non-histone proteins [55, 56]. While relatively little has been shown regarding the involvement of Setdb1 in SUDs, it has been identified in various neuropsychiatric and developmental disorders, such as Huntington’s disease [57], schizophrenia [58], and Rett syndrome [59]. Furthermore, Setdb1 is known to work with G9a-containing megacomplexes as a general transcriptional silencer via the methylation of H3K9 [60, 61]. Although direct connections between cocaine and Setdb1 are sparse, chronic cocaine exposure has been shown to dynamically regulate heterochromatic H3K9me3, one of the primary PTMs of Setdb1, in the NAc [50].

Consistent with G9a and Setdb1, Atf7ip was most highly expressed in the Cocaine-Craving cohort 21 days following the last cocaine exposure (**Fig. 3H**). Atf7ip is a multifunctional transcription factor associated with heterochromatin formation and stability, acting as a context dependent transcriptional regulator [62-64]. Evidence indicates that Atf7ip is necessary for Setdb1 stability and activity [64, 65], as Atf7ip and Setdb1 knockout cells exhibit nearly identical disruptions to global H3K9me3 [64]. Additionally, recent results suggest that the G9a/GLP complex can tri-methylate Atf7ip at an amino acid sequence similar to H3K9 [66], mediating its varying silencing activities, including interactions with Setdb1 [66]. In this context, Atf7ip may function as a recruiter for G9a and/or a binding-partner for Setdb1-containing silencing complexes.

The changes observed in G9a, Setdb1, Atf7ip, and Ash1l, along with increased H3K9me3 fluorescence intensity, suggest that one of the transcriptional regulatory mechanisms in VTA DA neurons in Cocaine-Craving mice could involve epigenetic repression mediated through H3K9. G9a and Setdb1 can both independently methylate H3K9 through various mechanisms, leading to transcriptional repression.

It is possible that propagation of H3K9 methylation may play a role driving transcriptional changes associated with cocaine craving. To explore this possibility, identifying the sequences repressed by H3K9me3 in VTA DA neurons during cocaine abstinence could yield potential drivers of cocaine relapse. To this end, this virus-based tagging method produces nuclei containing both RNA and chromatin [25], and as such, the transcriptional approaches described herein could also be coupled to novel chromatin profiling methods such as CUT&Tag, to examine specific loci exhibiting differential enrichment or depletion of H3K9me3 [67]. Such a combinatory approach could yield potential mutable targets associated with cocaine-craving, which may play a functional role in relapse to cocaine-seeking.

### Limitations

Complications with jugular catheterization, maintenance of catheter patency, and incomplete/off-target viral injection resulted in a 10-20% attrition rate. Further, limited numbers of VTA DA nuclei and RNA quantity per nucleus necessitated *in-house* cDNA library preparation, as opposed commercial alternatives (*See methods*). Lastly, as this study focused on method optimization and proof-of-principle, experiments were conducted using only male mice. While the methodology remains applicable to female mice, sex differences in drug metabolism [68], and potential variations in sequencing outcomes [27, 69] should be considered for future studies.

## Conclusion

We have refined an established nuclear labeling and capture method, paired with RNA-Seq in a complex behavioral paradigm, to provide distinct transcriptional profiles of VTA DA neurons during cocaine-taking and -craving. As an alternative to current molecular profiling methods, this platform enables targeted analysis of genetically defined neurons that can be adapted to study the role of dopamine signaling in other longitudinal SUD or neuropsychiatric disease models.

### Data and Code Availability

All code required for pre-processing of RNA-Sequencing data and differential gene expression analyses can be found at our Git repository: https://github.com/avm27/CocaineIVSA_RNASequencing_mDANeurons. All raw and necessary processed data used in this study can be found under GEO accession GSE277757.

## Acknowledgements

This work was supported by NIH grants K01DA045294 and DP1DA051828 (LMT), as well as a kind gift from the Shipley Foundation. FANS was performed with assistance from the Flow Cytometry Shared Resource (FCSR) of the Sylvester Comprehensive Cancer Center at the University of Miami, RRID: SCR022501, which is supported by NIH grant P30CA240139.

## Supplemental Results and Figures

**Supplemental Figure 1.**
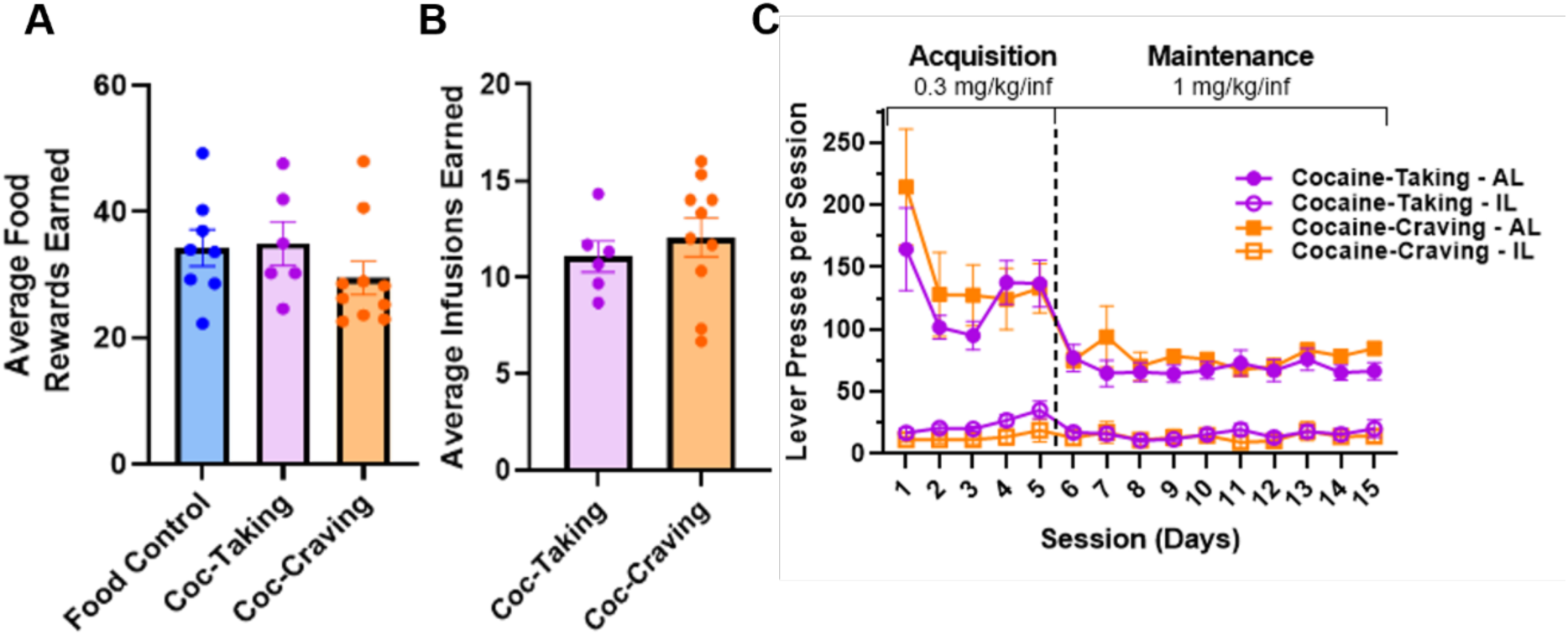
Comparisons of mouse behavior between selected experimental mouse cohorts. **A)** Average food rewards earned between selected mouse cohorts during the last 3 days of food training. Food Trained n=8; Cocaine-Taking n=6; Cocaine-Craving n=10 (one-way ANOVA, ns). **B)** Average cocaine infusions earned during the last 3 days of maintenance between cohorts during CIVSA (n=6-10/treatment group, unpaired t-test, ns). **C)** Active and Inactive lever presses between mice selected for Cocaine-Taking (purple) and Cocaine-Craving (orange) throughout CIVSA (two-way ANOVA, ns).

**Supplemental Figure 2.**
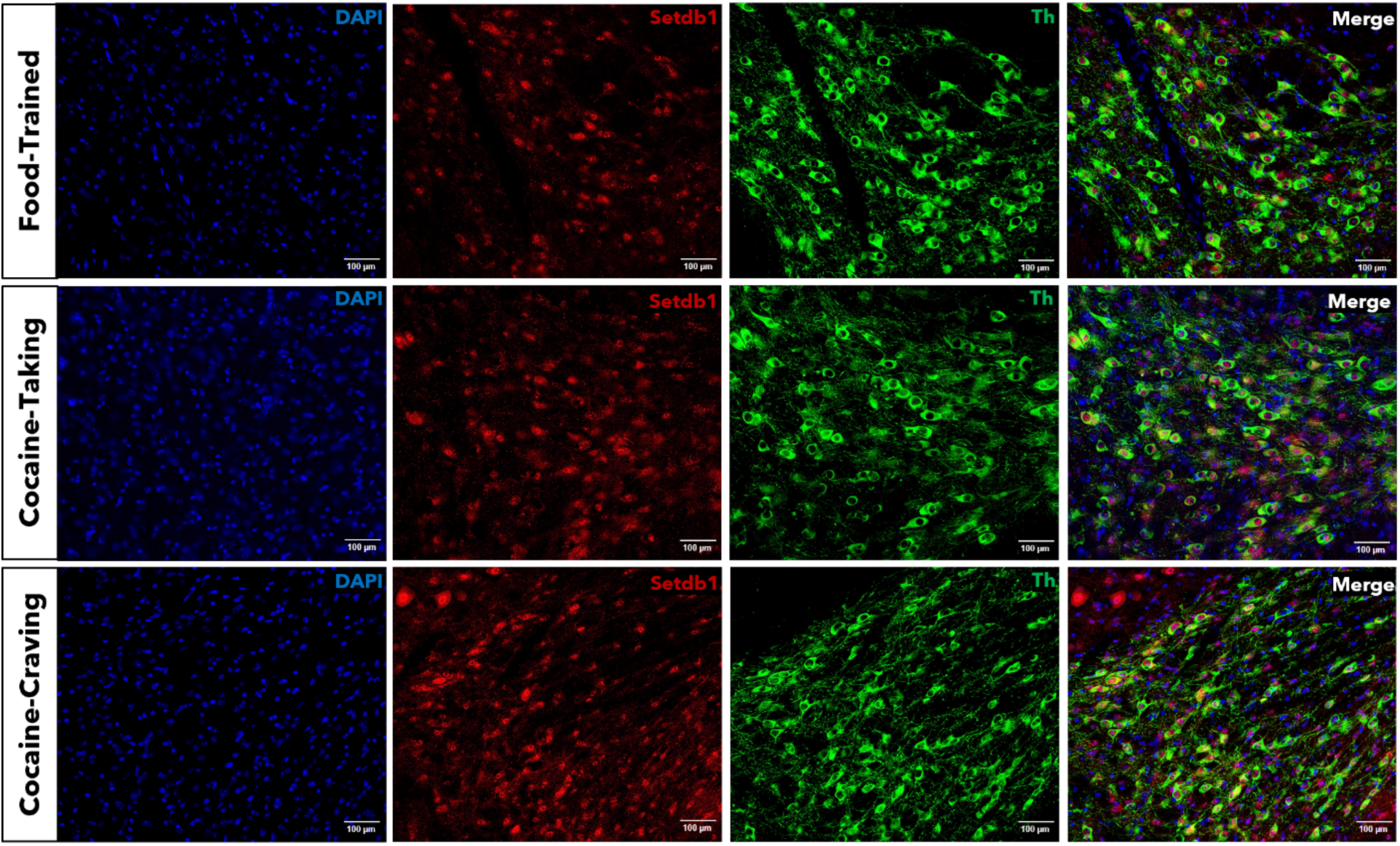
Immunolabeling of Setdb1 in VTA DA neurons throughout each phase of CIVSA. Representative IHC images of the VTA in Food Trained, Cocaine-Taking, and Cocaine-Craving in coronal mouse brain slices. Tyrosine hydroxylase (Th, green), SET domain bifurcated histone lysine methyltransferase 1 (Setdb1, red), and DAPI (blue) in 20x images of the VTA. Scale bar: 100µm.

**Supplemental Figure 3.**
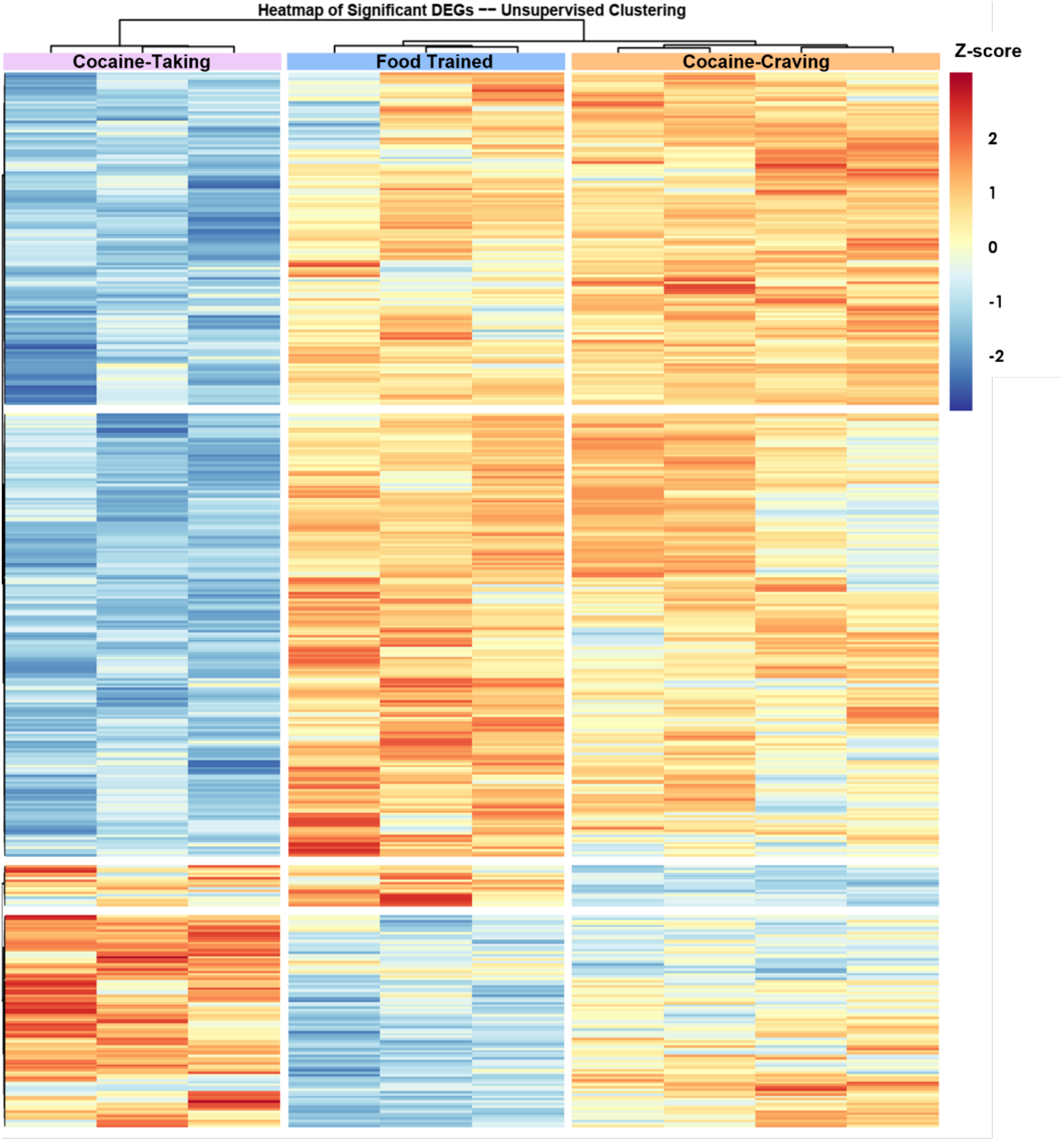
VTA DA neuron RNA independently cluster by treatment cohort. Unsupervised clustering heatmap of top 426 significant DEGs in HA+ nuclei between Food Trained, Cocaine-Taking, and Cocaine-Craving mice during CIVSA (padj < 0.05, L2FC ≥ 1.5 or L2FC ≤ -1.5, lfcSE ≤ 1).

